# Structural flexibility of the SARS-CoV-2 genome relevant to variation, replication, pathogenicity, and immune evasion

**DOI:** 10.1101/2021.12.20.473542

**Authors:** Roberto Patarca, William A. Haseltine

## Abstract

The SARS-CoV-2 pandemic continues to be driven by viral variants. Most research has focused on structural proteins and on site-specific mutations. Here, we describe recombination events involving genomic terminal sequences in SARS-CoV-2 and related viruses leading to structural rearrangements in terminal and coding regions and discuss their potential contributions to viral variation, replication, pathogenicity, and immune evasion.

## Introduction

Evolution of severe acute respiratory syndrome coronavirus 2 (SARS-CoV-2) variants involves both point mutations and recombination.^1–3^ The present analysis of the 5’- and 3’-untranslated regions (UTRs) of SARS-CoV-2 isolates from around the globe and from CoVs from bats in Thailand, Laos, and Great Britain revealed numerous previously undescribed duplications, inversions, and translocations within the genomic termini and into or from coding regions of the viral genome. Such variation may affect both clinical and epidemiological properties of the virus as the noncoding termini contain multiple regulatory sequences that are essential for viral gene transcription, translation, replication, and immune evasion.

## Materials and methods

Nucleotide and protein sequences were obtained from GenBank^®^ and the Global Initiative on Sharing All Influenza Data (GISAID) and searched and aligned using the Basic Alignment Search Tool (BLAST^®^, National Center for Biothechnology Information). RNA secondary structures were determined using forna, a force directed graph layout (ViennaRNA Web services).^4^

## Results

### Internal duplication and translocation of 5’-UTR segments in SARS-CoV-2 and related bat CoVs

The 5’-UTR of SARS-CoV-2 contains five stem-loop structures (SL) (Figure 1A).^5^ SL1 is required for the translation of viral mRNA.^6–8^ The function of SL2 is not well defined but based on similarity to other well studied CoVs it may be required for replication and transcription.^9,10^ SL3 encompasses the leader transcription regulatory sequence (TRS-L) required for messenger RNA synthesis.^11^ SL4 is involved in the synthesis of subgenomic RNA fragments, and the loops of SL5, which in SARS-CoV-2 include the initiation codon for the long 5’ open reading frames (ORF) 1a and 1ab, are likely involved in RNA packaging.^6^

**Figure 1.**
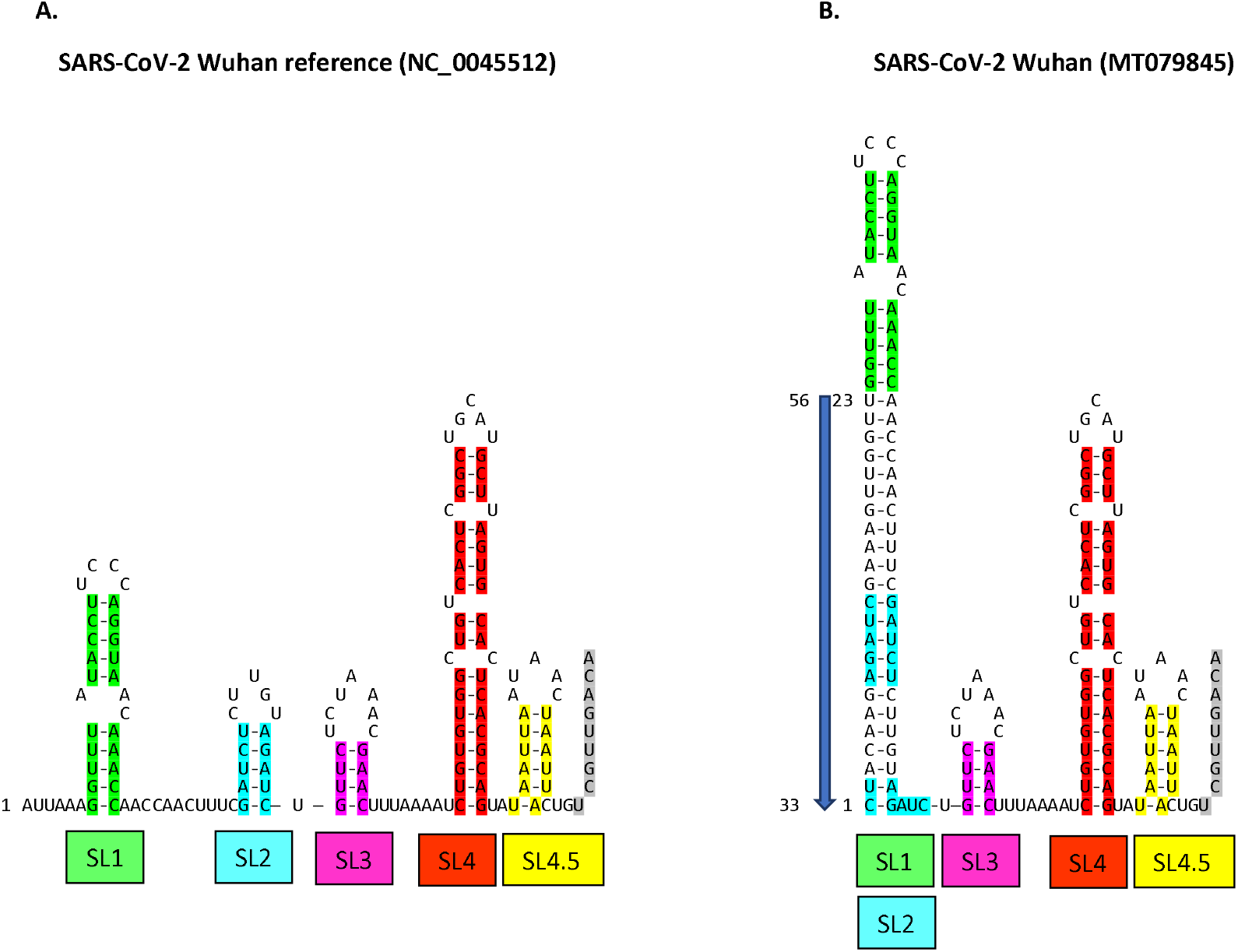
RNA secondary structures of: A. SARS-CoV-2 Wuhan reference (NC_0045512) and B. SARS-CoV-2 Wuhan (MT079845). Stem-loops (SL) 1 through 4.5 and beginning of SL5 (grey) are shown as predicted by forna algorithm and color coded. Insertion from minus-antisense strand (numbered according to position of nucleotides in Wuhan reference strain) is depicted by arrow.

The proximal ends of 5’-UTRs between the m^7^G cap and SL1 are reported to vary in length between one and six nucleotides.^6,12^ We noticed that the lengths of 5’-UTRs of SARS-CoV-2 and related bat CoVs differed in length and in distance between SL1 and the 5’-end well beyond a handful of nucleotides which led us to conduct an analysis of the 5’-UTR sequences. Our analysis revealed a striking variation in the length and composition of 5’ proximal sequences many of which exceed 100 nucleotides in length. Here we describe some representative samples of both the diversity and origin of the sequences.

Our search of GenBank and GISAID revealed the presence at the proximal end of the viral genome of exact inverted duplications of 5’-UTR sequences of various lengths (~20 to over 100 nucleotides). Figures 1B-3 depict examples derived from a long list of thousands of SARS-CoV-2 sequence entries from around the world including cruise ships and from that of the closely related Laotian bat CoV BANAL-20-236 and of the RacCS03 bat CoV from Thailand.^13^

**Figure 2.**
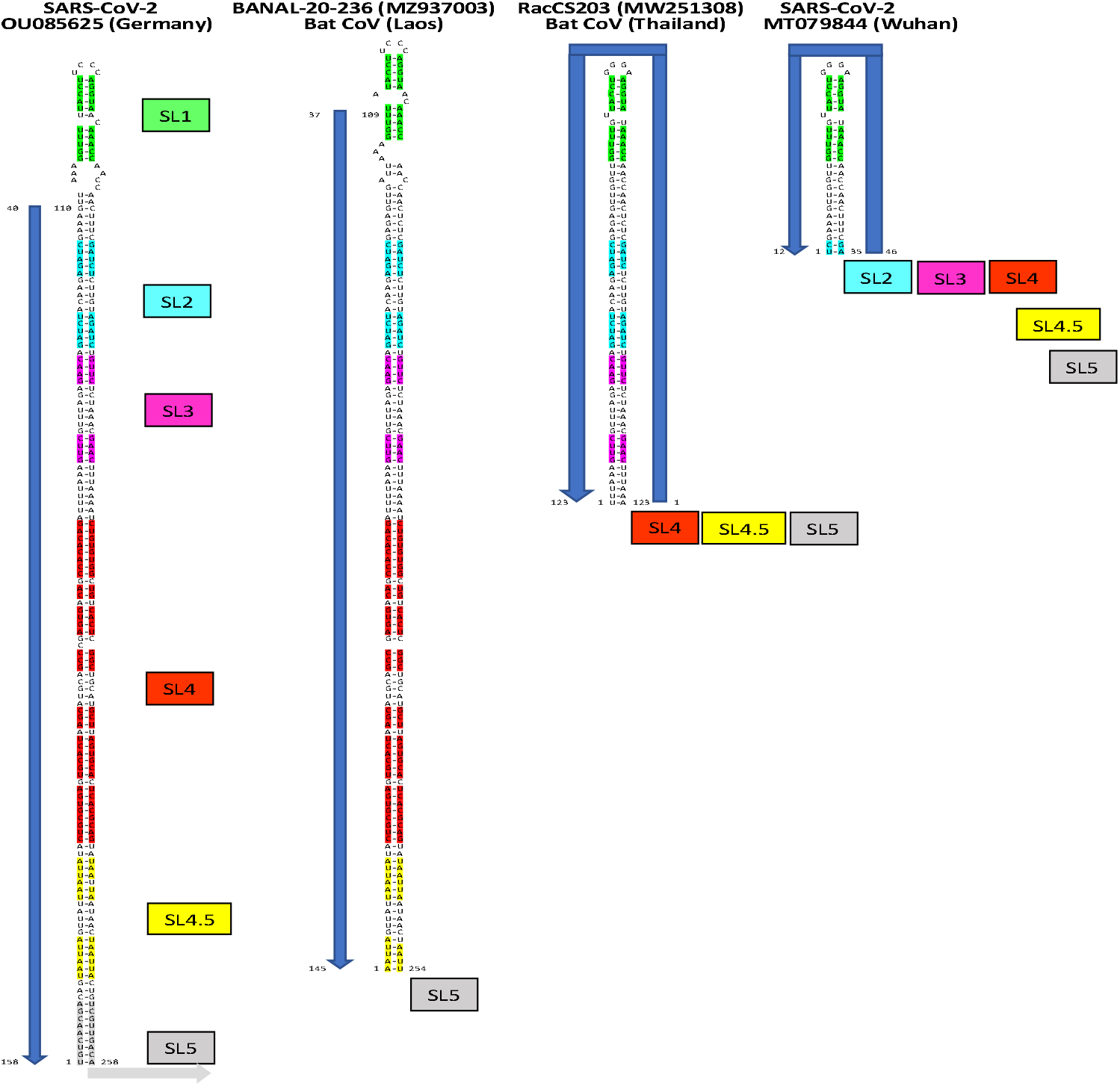
RNA secondary structures of: A. SARS-CoV-2 OU085625 (Germany), B. BANAL-20-236 bat CoV (MZ937003; Laos); C. RacCS203 bat CoV (MW251308; Thailand); and SARS-CoV-2 (MT079844; Wuhan). Stem-loops generated by minus-antisense strand insertions are shown with SL1-SL4.5 and beginning of SL5 color coded. Presence of stem-loops beyond first stem loop is shown by colored boxes. Notice that first two structures have a ‘CTTT’ in SL1 loop while the other 2 structures have ‘AGGG.’

**Figure 3.**
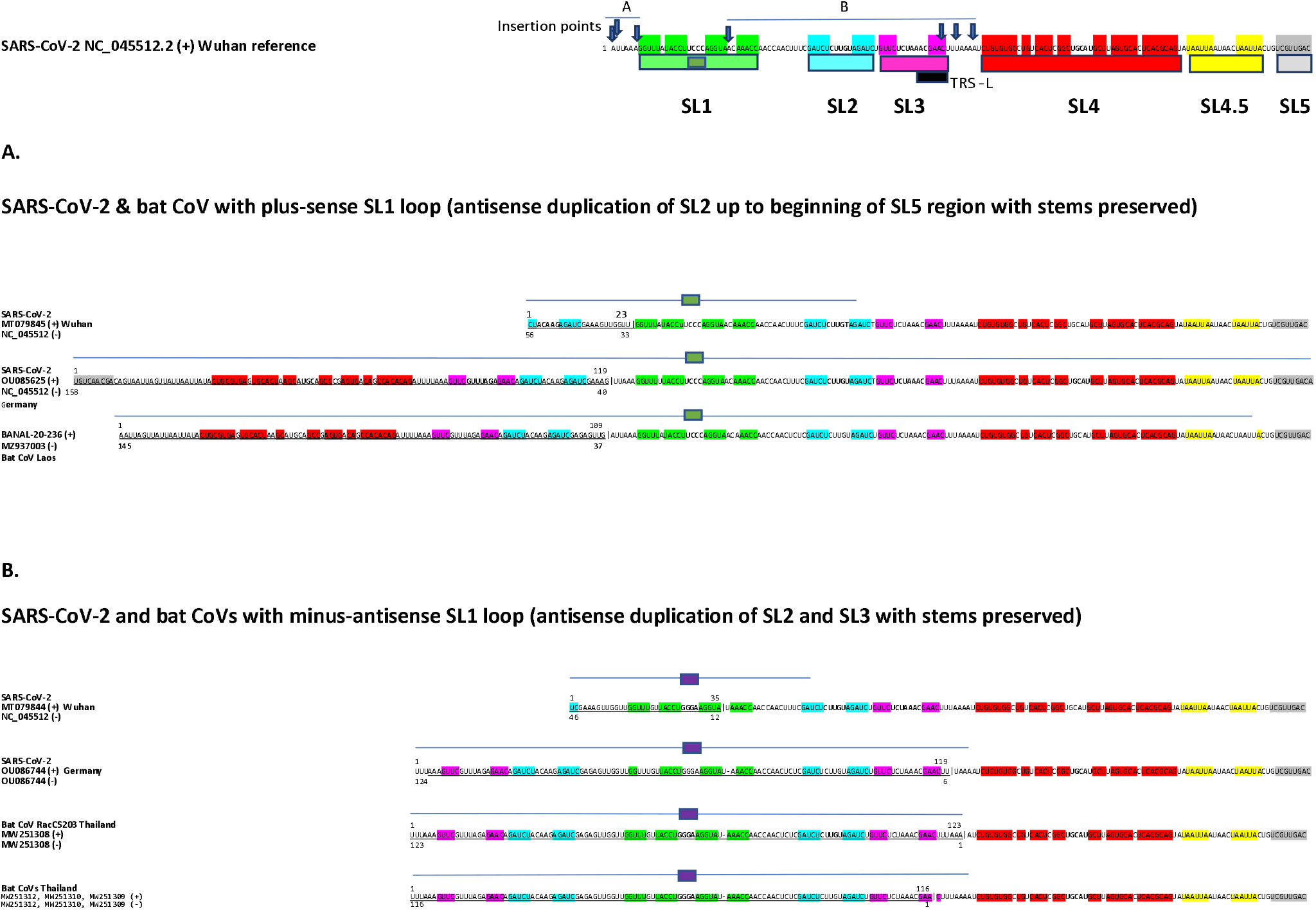
Summary of examples of primary structure of variants with minus-antisense strand insertions at the proximal end of the 5’-UTR. Primary structure of the SARS-CoV-2 reference strain is shown at the top with: SL1 through beginning of SL5 color coded; transcription regulatory sequence-leader (TRS-L) shown with a dark blue box; and the position of the insertions shown in panels A and B depicted with arrows. Panels A and B show examples of insertions resulting in SL1 loops with plus-sense (green box) and minus-antisense (purple box) loops, respectively. Insertions are underlined and span of minus-antisense strand is noted under the insertion sequences. Numbering of insertions is based on the indicated sequences used for their detection. Span of the long stem-loop that can be formed between minus-strand insertion and complementary plus-sense strand sequence is shown by a blue line. Note that the SL1 loop sits at the center of the hybrid long stem-loop.

Examination of secondary structures of the 5’UTRs yielded the surprising result of long stem-loop structures with shared structural features (Figs. 1-3). SL1 is conserved with subtle differences. The SL1 loop can be exactly that of the plus-sense strand (UCCC) or interestingly that of the minus-antisense strand (GGGA), while the stems of the SL1 structure are identical in sequence whether derived from a direct or inverted duplication, remarkably in the 5’ to 3’ direction.

The binding of nsp1 to SL1 is critical to viral mRNA translation. We infer that both forms of the SL1 sequence satisfy this requirement. This observation implies that the sequence of the stem is critical, and composition of the loop may vary in nsp1-SL1 interaction. This hypothesis requires testing.

We also note that the stems for SL2 to SL4.5 are conserved, and that their loops can collapse into the long stem. If the presence of single stranded loop sequences is required for the functions associated with SL2-SL4, there might be an equilibrium between the longer stem loop with SL1 up to SL4.5 stacked up in one hairpin structure and a structure with duplicated individual SLs 2 to 4.5. An equilibrium between two configurations might exist or helicases of viral and/or host origin may expose the loop regions which would be consistent with evidence garnered via a diversity of experimental means cited, while functionality of the long stem-loop structure might point to alternative mechanisms of translation and replication and even involvement of alternative host and viral proteins furthering our understanding of positive-sense stranded viruses.

The third shared feature is the presence of only one SL5 (Figure 1-3), which is consistent with SL5 folding into the same secondary structure independently of SL1-SL4.^6^

Further search of potential insertions of 5’-UTR sequences revealed duplication and translocation of 5’-UTR sequences not only within the 5’-UTR but also to coding regions of the viral genome (Fig. 4). We detected an insertion of a 27-nucleotide plus-sense strand segment of the 5’-UTR to the end of ORF8 gene in a SARS-CoV-2 variant isolated in Minnesota, USA and encoding an ORF8X protein with a modified carboxyl-terminus (5 last amino acids are replaced by 10 amino acids) (Figure 4A). It was previously noted that a longer overlapping 57-nucleotide segment of the 5’-UTR duplicated and translocated in place of an 882-nucleotide deletion within the coding portion of the viral genome of a SARS-CoV-2 variant isolated from 3 patients in Hong Kong with absent ORF7a, ORF7b, and ORF8 (lineage B.1.36.27) and encoding a C-terminally modified ORF6 product, termed ORF6X (Figure 4B).^14^

**Figure 4.**
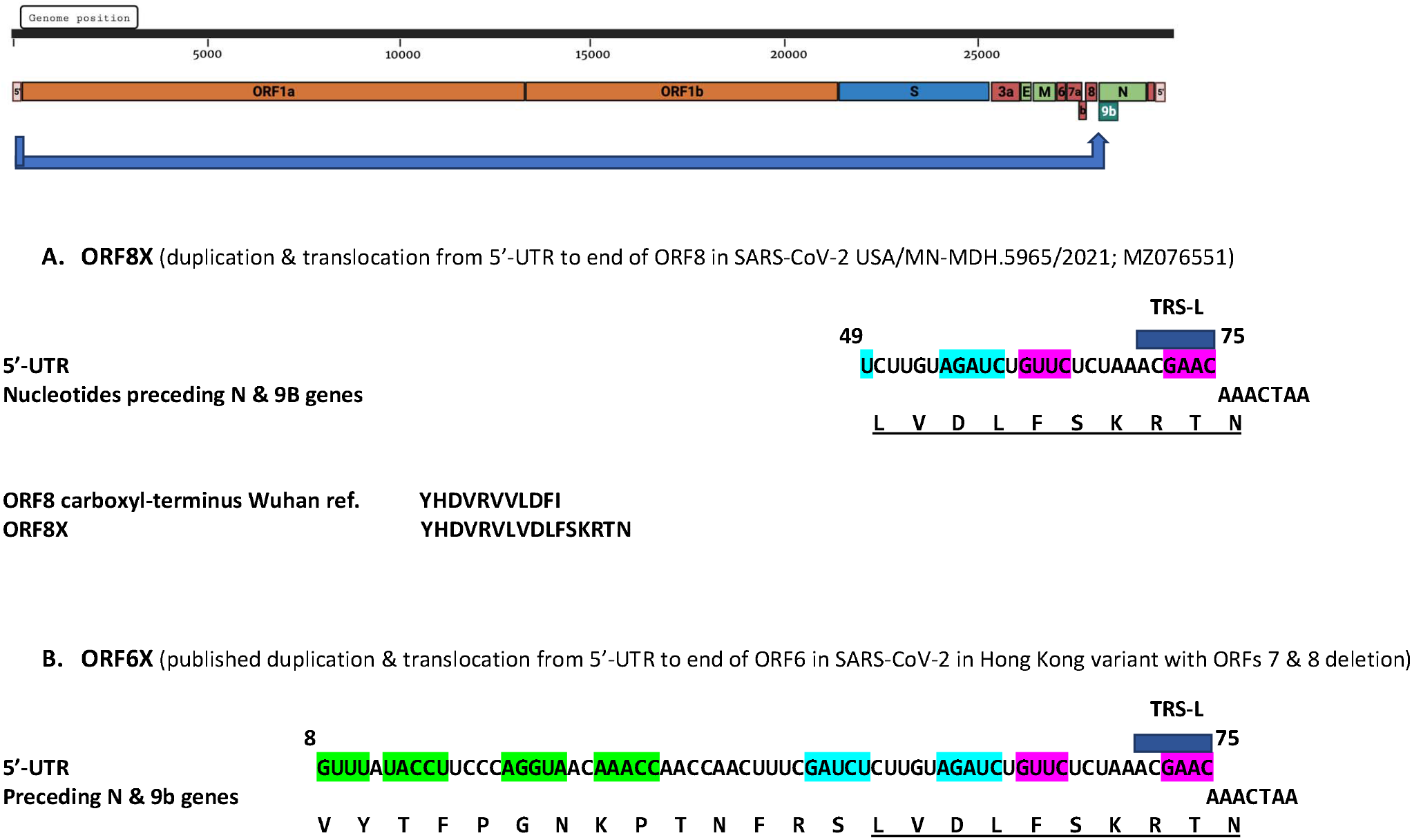
Duplication and translocation (long blue arrow in genome structure depiction) from 5’-UTR of SARS-CoV-2 variants to: A. end of coding sequence of ORF8 (SARS-CoV-2 USA/MN-MDH.5965/2021; MZ076551); and B. as published,^14^ end of coding sequence of ORF6 in a deletion variant lacking ORFs 7 and 8. Both insertions occur immediately before the beginning of N and ORF9b gene, and involve overlapping segments of 5’-UTR both ending in the TRS-L sequence in SL3. The last 5 amino acids of ORF8 are replaced with 10 amino acids encoded by the translocated 5’-UTR region and nucleotides and stop codon from the fusion region (underlined). The longer amino acid sequence translocated to the end of ORF6 is also shown.

We note that in both translocations, the end of the inserted 5’-UTR sequence corresponds to TRS-L that has been associated with spots with a higher frequency of recombination,^15^ and that insertion occurs at the same site immediately proximal to the nucleocapsid (N) and ORF9b genes, thereby bringing gene expression regulatory sequences to this location. These are the only insertions of 5’-UTR sequences into coronaviral genes characterized thus far.

### Duplication and translocation of viral gene segments to the 3’-UTRs of bat CoVs

We identified two instances of duplication, and/or inversion, and translocation of coding sequences at the end of the N gene and/or beginning of ORF10 to the distal end of the 3’-UTRs of two bat CoVs.

As shown in Figure 5, the 3’-UTR of the Laotian bat CoV BANAL-20-236 (MZ937003) has an additional 98-nucleotide segment at its distal end relative to that of SARS-CoV-2. An 88-nucleotide long subsegment in the corresponding minus-antisense strand is present in the plus-sense strand of the ORF10 region of SARS-CoV-2, BANAL-20 (Laos) and RhGB01 (England) bat CoVs, pangolin CoVs, and SARS-CoV immediately after the stop codon of the N gene and can form a stem-loop. Although the effects of said translocation remain to be determined, the loop of the minus-antisense strand of the ORF10 gene-associated stem-loop (GAAGAG) matches the six nucleotides in the octanucleotide (<u>GGAAGAGC</u>) that similarly protrudes in a loop in a hypervariable region of the murine hepatitis virus shown to be involved in viral pathogenicity.^16^ We note that the 88-nucleotide segment in the ORF10 region could also pair with the insertion sequence at the end of the 3’ UTR while retaining the loops present in the 3’ UTR.

**Figure 5.**
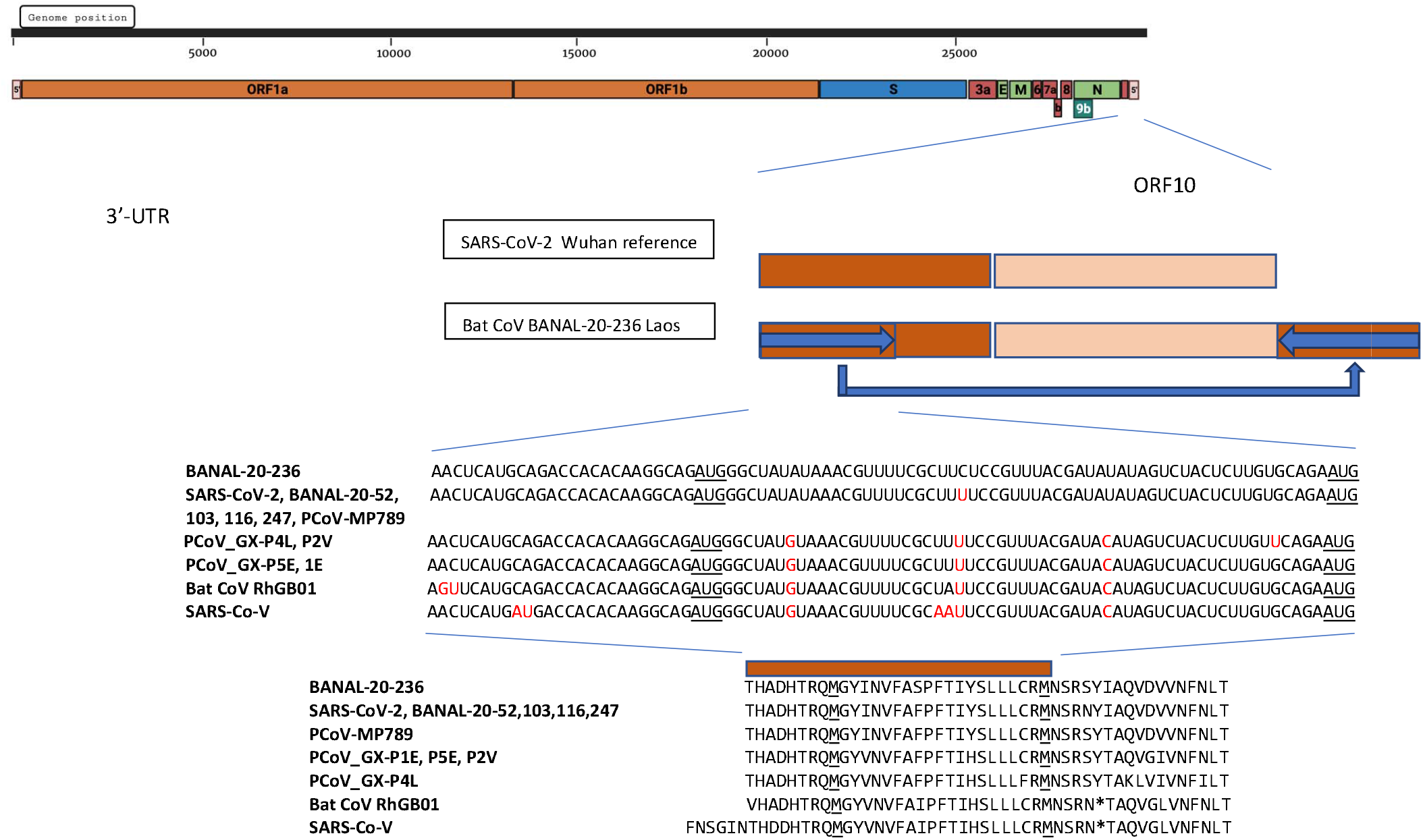
Duplication and translocation of minus-antisense strand from nucleotide segment in ORF10 to end of 3’-UTR in the BANAL 20-236 bat CoV (MZ937003). The nucleotide and protein sequences encoded by the ORF10 region are conserved among BANAL-20-236; SARS-CoV-2 (NC_0045512), BANAL-20-52 (MZ937000), BANAL-20-103 (MZ937001), BANAL-20-116 (MZ937002), BANAL-20-247 (MZ937004) bat CoVs (Laos), and pangolin (P)CoV-MP789; PCoV_GX-P4L (MT040333) and PCoV_GX-P2V (MT072864); PCoV_GX-P5E (MT040336) and PCoV_GX-P1E (MT040334); RhGB01 bat CoV (QYC92813; Thailand); and SARS-CoV (NC_004718).

In the case of the British bat CoV RhGB01 (MW719567), a 76-nucleotide plus-strand segment at the end of the N gene and beginning of the ORF10 gene is duplicated and translocated. Said sequence can fold into another stem loop (Figures 6 & 7), which as in the first case described can bring additional regulatory elements into the 3’-UTR.

**Figure 6.**
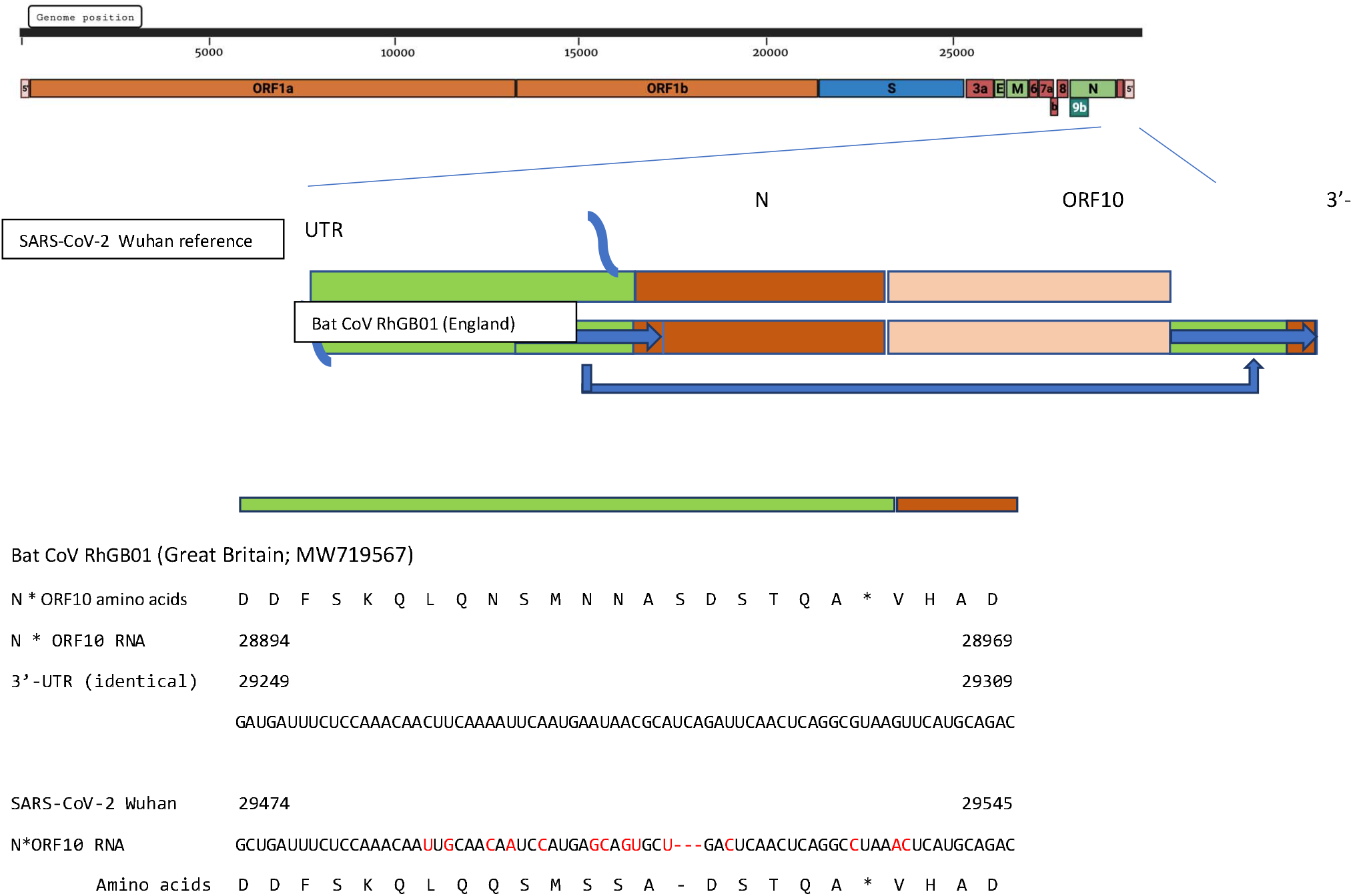
Duplication and translocation of plus-sense strand segment spanning end of N gene and beginning of ORF10 to end of 3’-UTR in the RhGB01 bat CoV (QYC92813, England). Alignments of the translocated nucleic acid and protein sequences of the bat CoV with the SARS-CoV-2 reference strain are shown.

**Figure 7.**
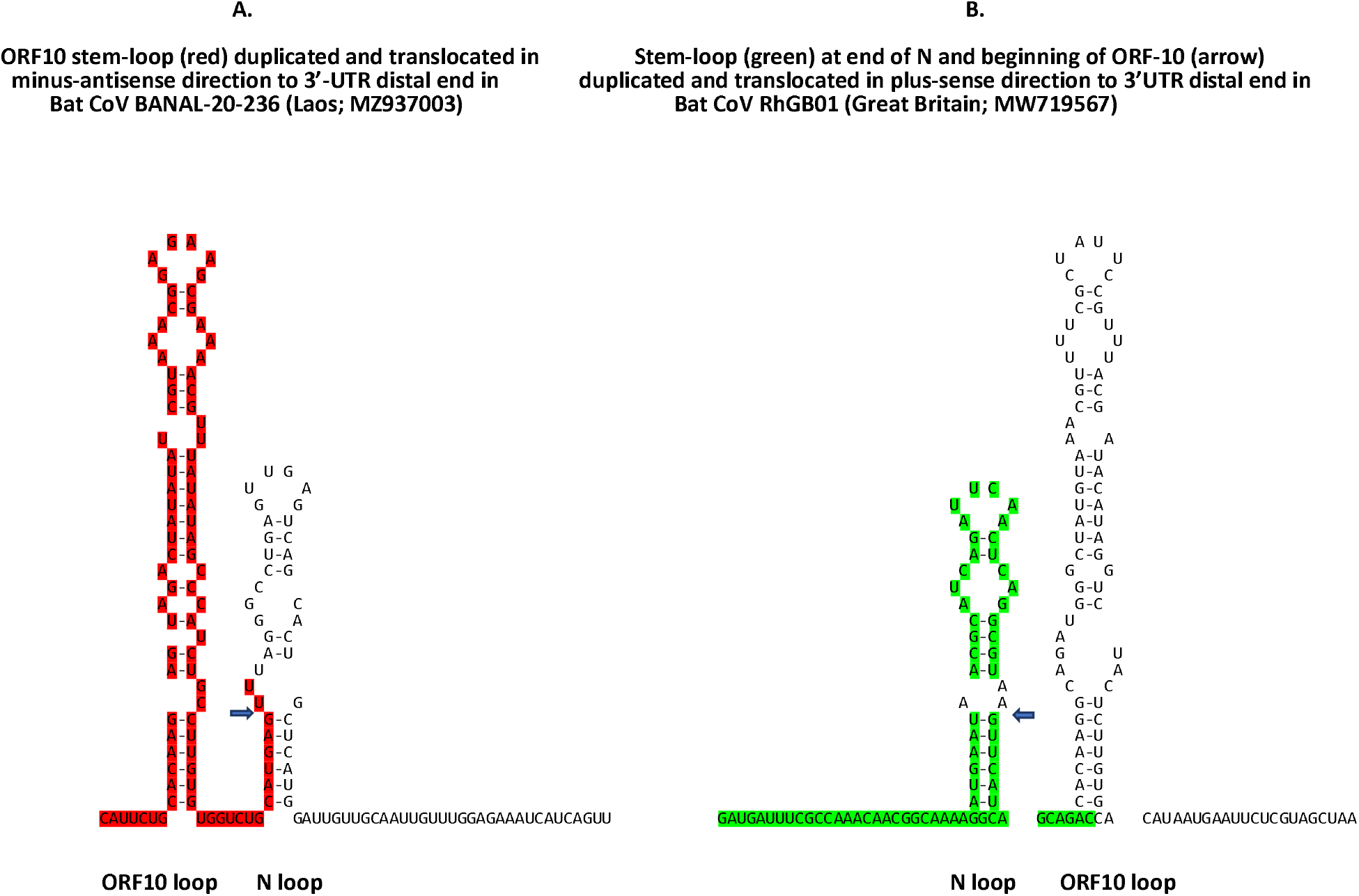
Stem-loops formed by the sequences translocated to the 3’-UTR shown in Figures 5 and 6. Panel A shows that derived from ORF10 region (present as shown in minus-antisense direction in 3’-UTR) in BANAL-20-236 bat CoV (MZ937003, Laos), and Panel B that derived from end of nucleocapsid N gene and translocated in the plus-sense direction to the end of the RhGB01 bat CoV (QYC92813, England) genome.

## Discussion

We here describe substantial duplications and rearrangements involving the 5’- and 3’-UTRs in addition to 5’-UTR-derived insertions into the coding region of the SARS-CoV-2 genome. These observations demonstrate the unanticipated flexibility of the terminal genomic regions of SARS-CoV-2. Most attention has focused on point mutations, and small insertions and deletions. As we consider the potential of future variants, we must be mindful of this structural flexibility as an inherent source of variation.

We considered the possibility that the variation described may be due to sequencing errors. However, the ubiquity of the variation in human and bat CoVs as well as its presence in replication-competent SARS-CoV-2 isolates as illustrated by those from Hong Kong with an anomalous ORF6 undermine this possibility. Moreover, we searched the GenBank database for similar insertions and found that at least 5,000 isolates from around the globe including cruise ships bear them.

Changes in length of viral genomic termini via insertions may serve an adaptive purpose and reflect a compensatory mechanism to address a common problem of linear genomes, namely, that they fray at both ends. Rather than acquiring segments from cellular mRNA as do influenza viruses, SARS-CoV-2 and related bat CoVs appear more parsimonious by deriving the insertions from their own genomes, and in the case of the 5’-UTR from the minus-antisense strand. Based on the observations of the present analysis there are restraints to this process. In the 5’-end, the SL1 loop in either plus-sense or minus-antisense orientation and the loops of SL5, both consistently present in one copy each, appear to be needed while the loops of SL2 to SL4.5 along with their conserved stems can become part of a long double-stranded stem. In the 3’-end, insertions at the end of the 3’-UTR that can form stem-loop structures may affect 3’-UTR-mediated regulation of gene expression, minus strand synthesis, and viral RNA stability and turnover as well as viral evolution. These observations call for a reexamination of the biochemical fundamentals of coronavirus replication and gene expression.

Correlation with infectivity, pathogenicity, and immune evasion data for the variants can provide insight into the effects of the genomic variations described here. For instance, the long 5’-UTR hairpin incorporating multiple SLs might render the virus more susceptible to 2’,5’-oligoadenylate synthetases-based immune response which relies on recognition of double stranded viral RNA.^17^ One could posit that other changes could be compensatory; for instance, the 3’-UTR stem-loop addition in BANAL-20-236 with a loop sequence that resembles that involved in pathogenicity of murine hepatitis virus may counteract potential untoward effects on immune susceptibility by the long stem-loop in its 5’-UTR.

Changes in viral genes secondary to translocation of 5’-UTR sequences may have other consequences. For instance, some diagnostic assays for SARS-CoV-2 infection target only accessory genes or proteins such as ORF8 and may be affected by ORF8 variants;^14,18^ however, the difference in the case here affects only the carboxyl-terminal end of ORF8. It remains to be determined how the changes described in viral genes like ORF6 and ORF8 affect their immune evasion-linked functions.

Recombination in coronaviruses occurs mainly via template switching, and transcription regulatory sequences (TRS) have been associated with recombination hotspots. Although canonical TRS-L dependent junctions at the proximal end of the subgenomic RNA encoding the N protein^19^ are most abundant, consistent with where the 5’-UTR insertions described here land, there are also non-TRS-dependent junctions towards the end of the N gene^19^ consistent with where the translocations to the 3’-UTRs in bat CoVs described here originate. In this respect, the nucleocapsid N protein in some coronaviruses might play a role in genome replication because it binds the 5’-UTR and intergenic sequences in mouse hepatitis virus RNA and is essential for infectivity of recombinant inflammatory bronchitis virus full-length transcripts.^20^

A caveat of the analysis performed here is that it relies on the quality of sequences deposited into GenBank and GISAID databases. Better algorithms to quality control and analyze sequences in a more automated fashion are warranted not only to better understand previously characterized variants but also to capture new ones more efficiently.

## Conclusion

The findings of this analysis of sequences available in databases illustrate the recombinatorial plasticity and coding potential of the UTRs of coronaviruses as contributors to viral evolution and the modification of existing viral genes. Whether the structural rearrangements described provide an advantage or disadvantage to the viral variants remains to be determined and correlations with viral infectivity, pathogenicity and immune evasion are warranted.

